# Distinct molecular signatures of fission predict mitochondrial degradation or proliferation

**DOI:** 10.1101/2020.11.11.372557

**Authors:** Tatjana Kleele, Timo Rey, Julius Winter, Sofia Zaganelli, Dora Mahecic, Hélène Perreten Lambert, Francesco Ruberto, Mohamed Nemir, Timothy Wai, Thierry Pedrazzini, Suliana Manley

## Abstract

Mitochondrial fission is a highly regulated process which, when disrupted, can alter metabolism, proliferation and apoptosis^1–3^. The downstream effects have implications for many diseases, from neurodegeneration^4–6^ to cardiovascular disease^7,8^ and cancer^9,10^. Key components of the fission machinery have been identified: constriction sites are initiated by the endoplasmic reticulum (ER)^11^ and actin^12^ before dynamin-related protein 1 (Drp1)^13^ is recruited to the outer mitochondrial membrane via adaptor proteins^14–17^, where it drives constriction and scission of the membrane^18^. In the life cycle of mitochondria, fission is important for the biogenesis of new mitochondria as well as the clearance of dysfunctional mitochondria via mitophagy^3,19^. Global regulation of fission on the cellular level is insufficient to explain how fate decisions are made at the single organelle level, so it is unknown how those dual functions arise, blocking progress in developing therapies that target mitochondrial activity. However, systematically studying mitochondrial division to uncover fate determinants is challenging, since fission is unpredictable, and mitochondrial morphology is extremely heterogeneous. Furthermore, because their ultrastructure lies below the diffraction limit, the dynamic organization of mitochondria and their interaction partners are hard to study at the single organelle level. We used live-cell structured illumination microscopy (SIM) and instant SIM^20^ for fast multi-colour acquisition of mitochondrial dynamics in Cos-7 cells and mouse cardiomyocytes. We analysed hundreds of fission events, and discovered two functionally and mechanistically distinct types of fission. Mitochondria divide peripherally to shed damaged material into smaller daughter mitochondria that subsequently undergo mitophagy, whereas healthy mitochondria proliferate via midzone division. Both types are Drp1-mediated, but they rely on different membrane adaptors to recruit Drp1, and ER and actin mediated pre-constriction is only involved in midzone fission.

## Positioning of mitochondrial fission site

At first glance, fission appears to occur randomly along the length axis of mitochondria (**Fig. 1a**). We recorded mitochondrial dynamics under physiological conditions at high temporal and spatial resolution, without pharmacological induction. We used live-cell SIM imaging of Cos-7 cells to precisely determine the position of fission for mitochondria of diverse shapes and lengths (**Fig. 1b**, **Supplementary Video 1**). Upon analyzing hundreds of spontaneous fission events, we discovered that their probability is not uniform; instead, they follow a bimodal distribution along a mitochondrion’s relative length (**Fig. 1c**). We term these either “peripheral” (positioned less than 25% of the total length from a tip) or “midzone” division (positioned within the central 50%). We obtained similar results when considering mitochondrial area instead of length (**Extended data Fig. 1a, b**), as expected because of the relatively constant mitochondrial diameter. This distribution was observed, independent of the total length of the dividing mitochondria (**Fig. 1d, Extended Data Fig. 1c**). As a consequence of this bimodality, smaller daughter mitochondria derived from peripheral divisions have a relatively narrow length distribution (1-2 μm) (**Fig. 1e**). Mitochondria labelled with either inner or outer membrane markers revealed a similar bimodal distribution, confirming fission was complete (**Extended Data Fig. 1d**). To test whether our observations are generally reflective of mitochondrial fission in mammalian cells, we repeated our measurements in postnatal mouse cardiomyocytes, a different cell type and species (**Fig. 1f**, **Supplementary Video 2**). Indeed, we recovered a bimodal distribution with mitochondria dividing either in the midzone or peripherally (**Fig. 1g**); thus, we wondered whether these two geometrically distinct fission types signified underlying physiological or functional differences.

**Fig. 1.**
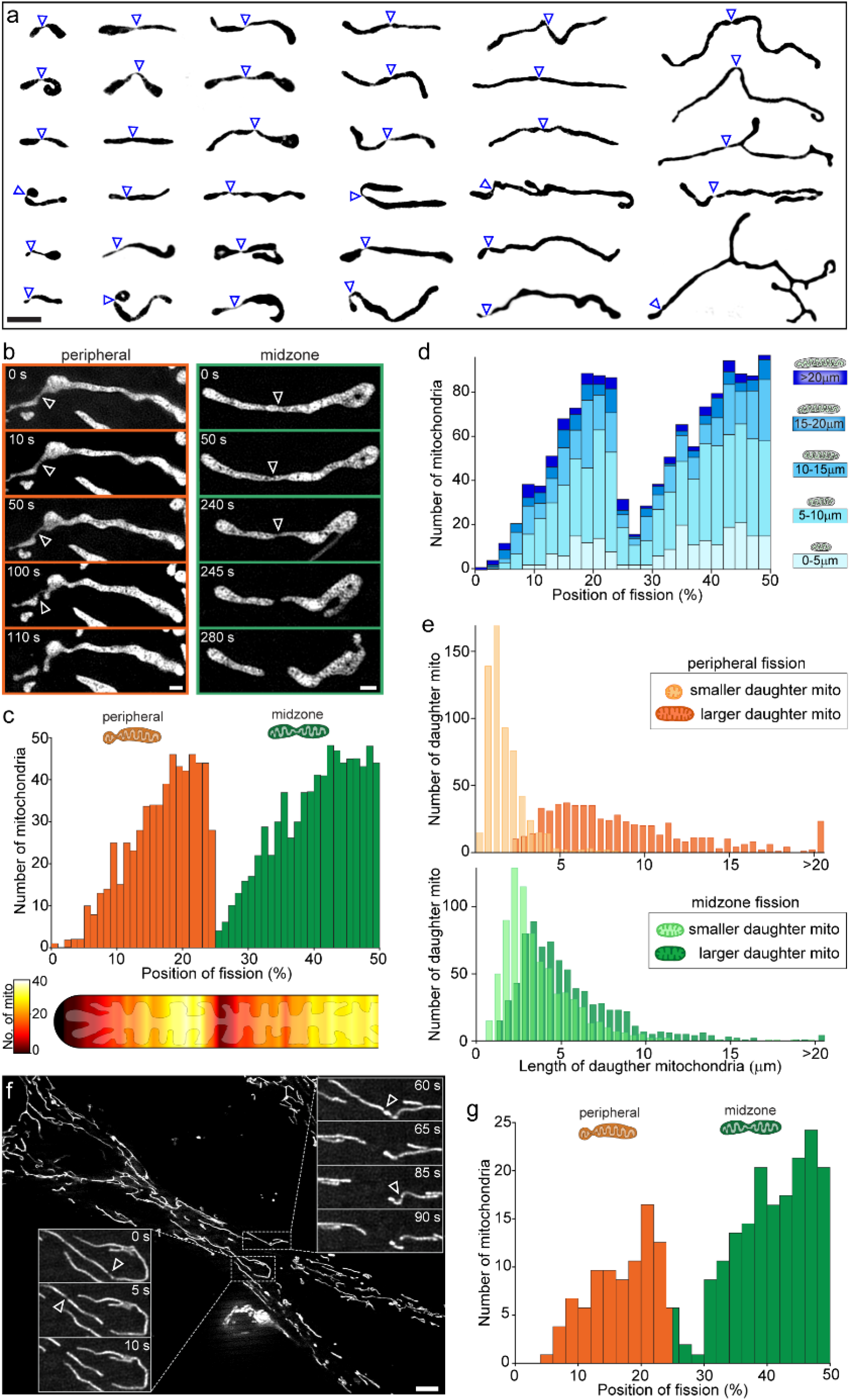
Mitochondrial fission sites are bimodally positioned. **a**, Gallery of examples of mitochondria in Cos-7 cells one frame before division from time-lapse SIM movies, represented as binary images. **b**, Time-lapse sequences from SIM movies showing a peripheral (left) and a midzone (right) fission of mitochondria labelled with Mitotracker green. **c**, Histogram of the relative position of fission along the length axis of Cos-7 mitochondria (n=1393 fissions pooled from different data sets) shows a bimodal distribution, with fissions more likely to occur near the tip (orange, 0-25% bin; ‘peripheral’) or near the center (green, 25-50% bin; ‘midzone’). The heat map (below) illustrates the probability of fission along a mitochondrion. **d**, Stacked histogram of the relative position of fission for different bins grouped by the total length of the dividing mitochondria (replot from data in **c**). **e**, Length distribution of the smaller (light color) and larger (dark color) daughter mitochondria arising from peripheral (top, orange) and midzone (bottom, green) fissions. **f**, Live-cell iSIM image of mitochondria (Mitotracker) in primary, postnatal mouse cardiomyocytes. Insets show two time-lapse sequences of fission events from indicated areas (box). **g**, Histogram of the relative position of fission in cardiomyocyte mitochondria (n=380 fissions), represented as in **c**. Scale bars are 0.5 μm in **b** and 10 μm in **f**. Fission sites are indicated by arrowheads.

## Signs of stress and dysfunction precede peripheral fission

Mitochondria are a hub for different metabolic functions, characterized by distinct physiological and biochemical properties. The potential across the inner membrane drives oxidative phosphorylation, creating a pH difference between matrix and intermembrane spaces. Production of reactive oxygen species (ROS), a toxic byproduct of oxidative phosphorylation, can lead to mitochondrial damage, often accompanied by loss of membrane potential and release of Ca^2+^ and cytochrome c^21^. We investigated the physiological states of mitochondria preceding fission with fluorescent sensors and live-cell SIM (**Fig. 2**). We found mitochondrial membrane potential as reported by the dye TMRE was reduced prior to fission in small peripheral daughter mitochondria (**Fig. 2a, b**, **Supplementary Video 3**) when compared with corresponding large daughter mitochondria or non-dividing mitochondria. In contrast, no differences were observed between daughter mitochondria from midzone fissions. Similarly, the genetically encoded pH sensor SypHer^22^ reported a reduced matrix pH in small daughter mitochondria prior to fission (**Fig. 2c, d**, **Supplementary Video 4**). The lower fluorescence intensities of small daughter mitochondria seen for TMRE and mito-SypHer were not observed for mito-GFP, and thus cannot be explained by their reduced volume or uptake (**Extended Data Fig. 2a, b**). Furthermore, the smallest mitochondria derived from midzone fission are smaller than the largest peripheral daughter mitochondria, underlining that the differences we detect cannot be explained by size (**Extended data Fig. 2c-i**).

**Fig. 2.**
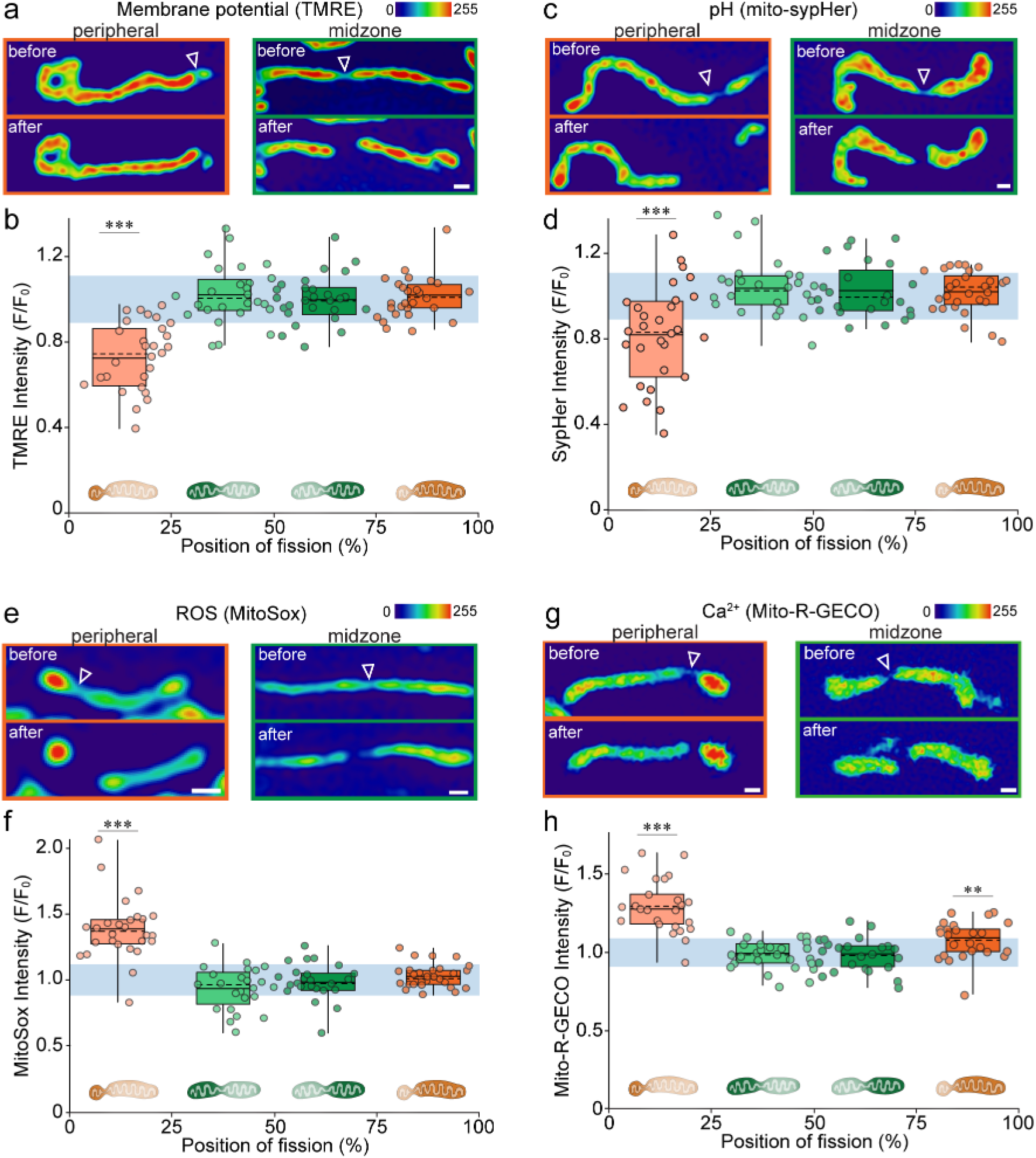
Peripherally dividing mitochondria display signs of stress and dysfunction. **a**, Mitochondrial membrane potential before and after example peripheral (left) and midzone (right) fissions taken from time-lapse SIM movies of TMRE stained Cos-7 mitochondria. **b**, Normalized TMRE intensity depending on the relative position of fission measured in mitochondria immediately before fission. Circles indicate individual measurements and average values of binned groups (0-25%, 25-50%, 50-75% and 75-100%;) are represented as box plots (box= 25-75 percentile; bars= min/max values; dotted line=median; solid line= average; n=54 fissions). Light blue area shows the average TMRE intensity in non-dividing mitochondria (±SD). **c**, Matrix pH in dividing mitochondria taken from SIM movies of mito-SypHer transfected Cos-7 cells. **d**, Normalized mito-SypHer intensity depending on the relative fission position represented as in **b**. Light blue area shows the average mito-SypHer intensity in non-dividing mitochondria (±SD; n≥ 25 fissions for each group). **e** Examples of MitoSox labelled Cos-7 mitochondria dividing in the periphery (left) and midzone (right) taken from ratiometric measurements. **f**, Normalized MitoSox intensity depending on the position of fission represented as in (**b)**. Light blue area shows the average MitoSox ratio in non-dividing mitochondria (±SD; n≥ 24 fissions for each group. **g**, Images of mito-R-Geco transfected Cos-7 mitochondria immediately before peripheral (left) and midzone (right) fission. **h**, Normalized mito-R-Geco intensity depending on the position of fission, represented as in (**b)**. Light blue area shows the average mito-R-Geco intensity in non-dividing mitochondria (±SD; n≥ 31 fissions for each group). Statistical significance calculated by two-tailed t-test for normally distributed populations and Mann Whitney U test for non-normally distributed populations; *P< 0.05, **P<0.01, ***P<0.001. Scale bars are 0.5 μm. Fission sites are indicated by arrowheads.

Another indicator of mitochondrial dysfunction is the accumulation of ROS, usually eliminated by anti-oxidative enzymes^23,24^. Consistent with the reduced membrane potential and pH we observed in small peripheral daughter mitochondria, ROS levels measured by the fluorogenic dye MitoSox (**Fig. 2e, f**) and the genetically encoded ROS sensor mito-Grx1-roGFP (**Extended Data Fig. 2j, k**) were elevated compared to non-dividing mitochondria or daughters from midzone fissions. To test the effect of ROS scavengers on peripheral fissions, we treated cells with 500 nM MitoQ^25^ and found that peripheral fission rates are decreased, while there was no effect on midzone fission rates (**Extended Data Fig. 2f**). Finally, to examine mitochondrial Ca^2+^ levels, we used the genetically encoded sensor mito-R-Geco^26^ (**Fig. 2g, h**, **Supplementary Video 5**) and Cepia3-mt^27^ (**Extended Data Fig. 2l, m)**. Mitochondrial Ca^2+^ plays important roles in cell survival and death by balancing homeostasis^28,29^. We observed Ca^2+^ levels increased significantly in small peripheral daughter mitochondria, and mildly in large daughter mitochondria, compared with midzone or non-dividing mitochondria. Thus, we found no differences in the physiological states of midzone fissions before or after fission, whereas peripheral fission is preceded by a rise Ca^2+^ and in ROS, as well as reduced membrane potential and pH near the tip of the mitochondrion that produces the small daughter organelle.

## Midzone fission serves biogenesis while peripheral fission is followed by mitophagy

We next investigated the functional outcomes of midzone and peripheral fissions. Since the physiology of individual mitochondria showed signs of stress and damage upstream of peripheral fission, we hypothesized that it could be linked to degradation, whereas midzone fission could serve biogenesis. To test this, we analyzed the redistribution of mitochondrial DNA (mtDNA) in Cos-7 cells stained with the vital dye PicoGreen (**Fig. 3a**, **Extended Data Fig. 3a)**. On average, there was no significant difference in the total number of mtDNA foci (nucleoids) in peripherally versus centrally dividing mitochondria. However, we observe a bimodal distribution of nucleoids in smaller daughter mitochondria from peripheral fissions, with 32% containing no nucleoids (versus 3% in the other daughter mitochondria). Mitochondria lacking mtDNA also had diminished membrane potential compared to those containing mtDNA (**Extended Data Fig. 3b**). We observed similar results for mitochondrial RNA granules (MRGs, **Extended Data Fig. 3c-e**), which are composed of mtRNA and RNA processing proteins^30^. It has been proposed that ER-mitochondrial contacts coordinate licensing of mtDNA replication, thereby coordinating synthesis with division^31^. Daughter mitochondria arising from midzone fissions contained a high number of replicating (Mito-Twinkle^32^ positive) nucleoids compared to non-dividing mitochondria, consistent with a role in mitochondrial proliferation (**Fig. 3b, Extended Data Fig. 3f**). In contrast, 75% of smaller peripheral daughter mitochondria contained no Twinkle foci. Since peripheral fission might also remove damaged mtDNA, we exposed cells to UV light (**Fig. 3c**), and labelled newly synthesized RNA in treated versus control cells with Bromo-Uridine (BrU). Indeed, mitochondria after UV treatment have no or very few BrU foci, indicating disruption of transcription, presumably due to mtDNA damage (**Extended Data Fig. 3g**). Consistent with our hypothesis, the prevalence of nucleoids increased in small peripheral daughter mitochondria (82% versus 68% without UV irradiation).

**Fig. 3.**
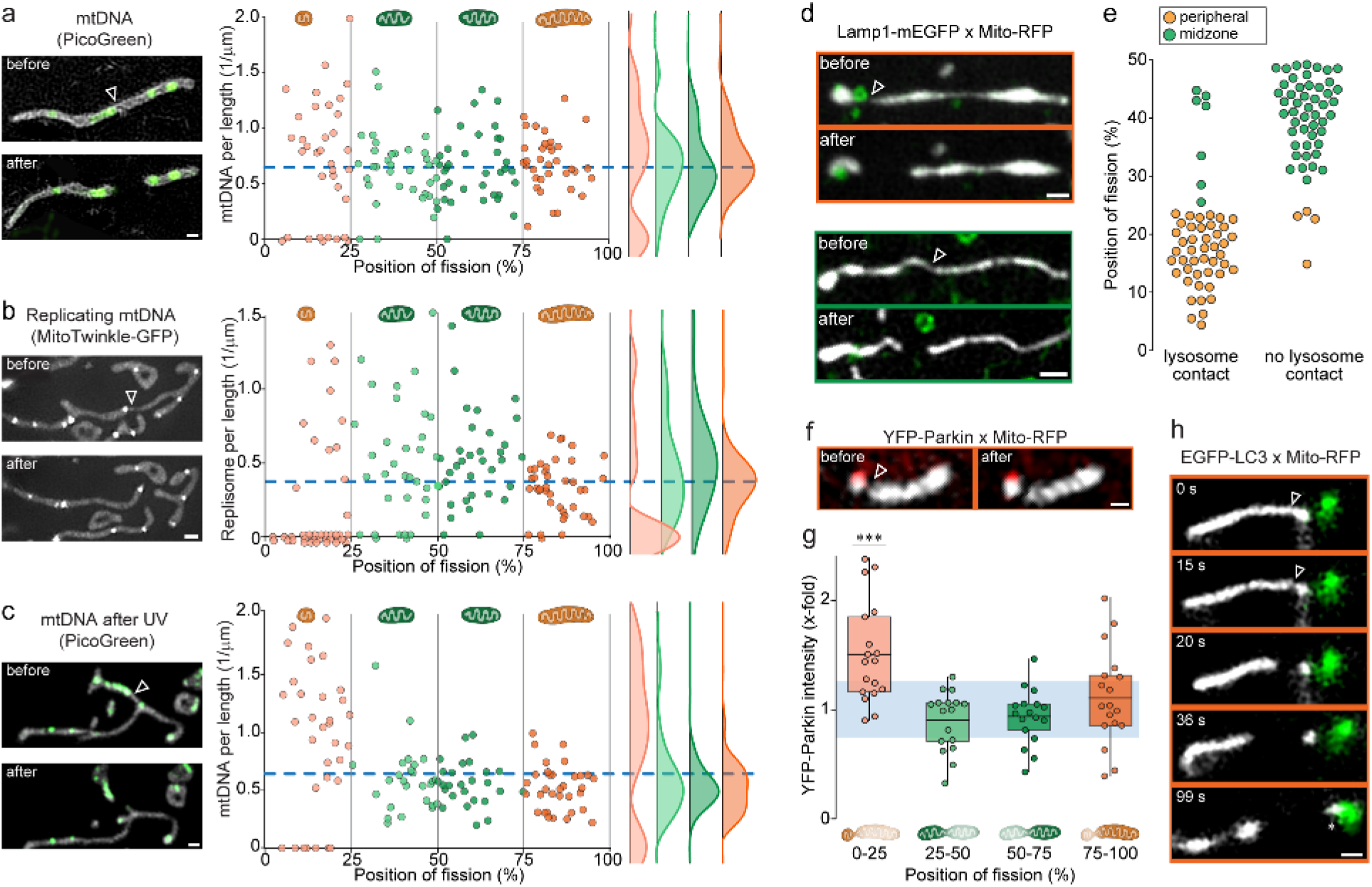
Midzone division is coupled with mtDNA replication and peripheral division precedes mitophagy. **a**, SIM images of Cos-7 mitochondria (mito-RFP, greyscale) and mtDNA (PicoGreen, green) before and after fission (left). Quantification of mtDNA foci per μm length depending on the fission position (right). Circles are individual measurements. Curves show frequency distributions of binned groups (0-25%, 25-50%, 50-75%, 75-100%). Blue line displays average mtDNA in non-dividing mitochondria (n=78 fissions). **b**, SIM images and quantification of replicating mtDNA (mito-Twinkle-GFP) per μm length depending on the fission position, represented as in **a**(n= 76 fissions). **c**, SIM images and quantification of mtDNA foci depending on the fission position after UV irradiation represented as in **a**(n= 64 fissions). **d**, SIM images of mitochondria (mito-RFP, greyscale) and lysosomes (Lamp1-mEGFP, green) before and after peripheral and midzone fission. **e**, Position of fission in divisions showing a contact or no contact with lysosomes prior to fission (n= 51 fissions). **f**, SIM images before and after fission in Cos-7 cells transfected with YFP-Parkin (red) and mito-RFP (greyscale). **g**, Normalized pre-fission YFP-Parkin intensity depending on the fission position. Circles indicate individual measurements and average values of binned groups (0-25%, 25-50%, 50-75%, 75-100%;) are represented as box plots (box= 25-75 percentile; bars= min/max values; solid line= average; n=34). Light blue area shows average YFP-Parkin intensity in non-dividing mitochondria (±SD). **h**, Time-lapse SIM sequence of Cos-7 mitochondria (mito-RFP, grey) and autophagosomes (EGFP-LC3B, green), where the small daughter mitochondrion from a peripheral fission is being taken up by an autophagosome (asterisk). Statistical significance calculated by two-tailed t-test for normally distributed and Mann-Whitney U test for non-normally distributed populations; ***P<0.001. Scale bars are 0.5 μm. Fission sites indicated by white arrowheads.

To test our hypothesis of the degradative function of peripheral fission, we followed the fate of resulting daughter mitochondria. Previous studies reported lysosome-mitochondria contacts prior to fission^33^, which we observed in 92% of peripheral fissions, using Lamp1-mEGFP as a marker, compared with only 13% of midzone fissions (**Fig. 3d, e**, **Supplementary Video 6**). At peripheral fission sites, we also frequently found mitochondrial-derived vesicles (MDVs), formed from the outer membrane and known to be targeted to late endosomes for degradation^34^ (**Extended Data Fig. 6c, d**). Mitochondria undergoing peripheral fission also accumulated YFP-Parkin on their membranes (**Fig. 3f, g**). These results link peripheral fission with mitophagy, which is the turnover of excess or dysfunctional mitochondria by autophagy, regulated by PINK1 and Parkin^35^. In some cases, we could track the small daughter mitochondria derived from peripheral fissions over several minutes and observe their uptake by autophagosomes (**Fig. 3h**, **Supplementary Video 7**). Thus, peripheral fission is implicated in quality control and initiation of organelle clearance by mitophagy. To dissect the sequence of events leading to fission, we analyzed the timing of changes in mitochondrial physiology (**Fig. 2**) and recruitment of the fission and autophagic machinery. This revealed that a drop in membrane potential, rise in Ca^2+^, and recruitment of the autophagic machinery all precede peripheral fission (**Extended Data Fig. 4**).

Previous studies established a paradigm whereby dividing mitochondria either re-fuse with the mitochondrial network, or are isolated and undergo mitophagy^3^. Therefore, we followed the fate of daughter mitochondria after division. We found that small peripheral daughter mitochondria are excluded from further fusions or divisions in both Cos-7 cells (**Fig. 4a**) and primary mouse cardiomyocytes (**Extended Data Fig. 5a**), while the remaining daughter mitochondria regularly undergo further fission or fusion events.

**Fig. 4.**
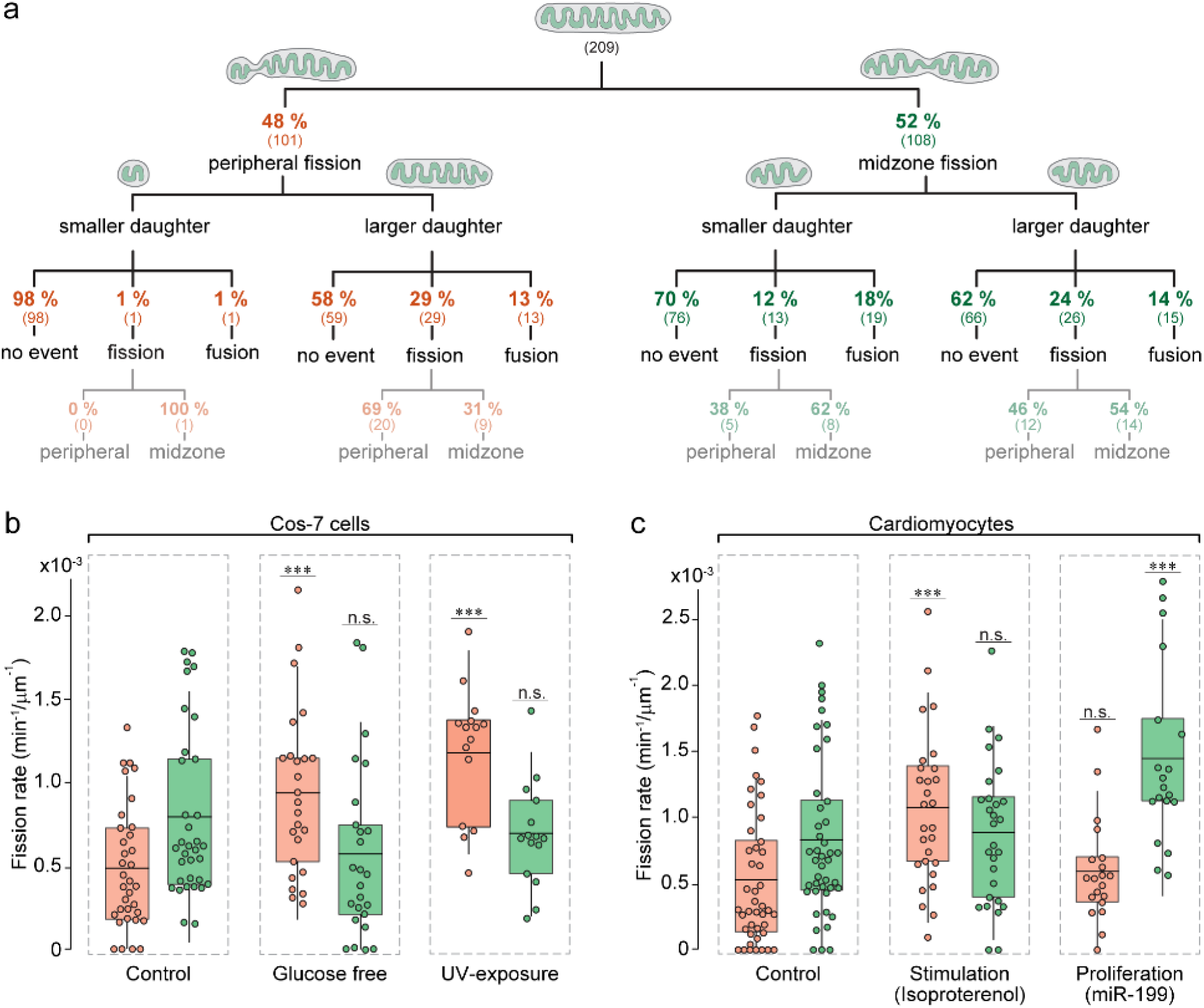
Peripheral and midzone fissions can be adapted independently depending on the cellular context and interact differently with the mitochondrial network. **a**, Schema depicting the fate (”no event”, another “fission” or “fusion”) of each daughter mitochondrion from peripheral and midzone fissions after the initial division in Cos-7 cells. Only mitochondria that could be traced for more than 100 seconds post-fission were included. Numbers in parenthesis are total numbers of events. **b**, Quantification of the peripheral (orange) and midzone (green) fission rates (number of fissions per minute per μm length) in control Cos-7 cells and cells starved in glucose-free medium for 48 hours. **c**, Fission rates in postnatal mouse cardiomyocytes under control conditions, after 48 of stimulation with Isoproterenol or after inducing proliferation with a miR-199 mimic (box= 25-75 percentile; bars= min/max values; solid line= average; n≥ 20 cells for each group). Statistical significance calculated by two-tailed t-test for normally distributed populations and Mann Whitney U test for non-normally distributed populations; n.s. P>0.05, ***P<0.001.

## Peripheral and midzone fissions are modulated independently depending on the cellular context

We wondered whether cells would be able to modulate each fission type independently, in a context-dependent manner (**Fig. 4b, c**). Thus, we subjected Cos-7 cells to metabolic stress by growing them in glucose-free (galactose supplemented) media^36^ for 48 hours. We observed that the rate of peripheral fissions per cell increased, whereas the rate of centrally dividing mitochondria remained constant (**Fig. 4b, Extended Data Fig. 5b**). A similar trend emerged when we exposed Cos-7 cells to UV light to induce mtDNA damage. We also tested the effects of increased energy demand and hence oxidative stress on primary mouse cardiomyocytes, by treatment with isoproterenol, a non-selective β-adrenergic receptor agonist increasing contractility and inducing hypertrophy. We imaged cells after 48 hours of constant stimulation and found increased rates of peripheral fissions compared to non-treated cells, while the rate of midzone fissions remained constant (**Fig. 4c, Extended Data Fig. 5c**). In contrast, when cells proliferate, our model predicts an upregulation of midzone fissions. To test this, we treated cardiomyocytes with a miR-199 mimic^37^; indeed, we found increased rates of midzone fissions, whereas peripheral fission rates remain unchanged (**Fig. 4c, Extended Data Fig. 5d**). Thus, cellular stress and high energy demands, associated with oxidative damage, increase the rate of peripheral fissions, whereas cell proliferation, which requires biogenesis of new mitochondria, increases the rate of midzone fissions.

## Peripheral and midzone fissions are mediated by different molecular machineries

The differences in physiology and fate of mitochondria derived from these two types of fissions prompted us to investigate the molecular players involved. Previous studies report that mitochondrial-ER contacts^11^, in coordination with actin polymerization^38^, define division sites and trigger mitochondrial DNA replication upstream of Drp1, and sometimes Dyn2^39–41^ assembly. All fissions we observed were spontaneous, and mediated by Drp1 assembly in both midzone and peripheral cases (**Extended Data Fig. 6a, b**). In two-color measurements of the endoplasmic reticulum (ER) and mitochondria, we found that midzone fission sites consistently contacted the ER prior to fission, but most peripheral fission sites did not (**Fig. 5a, b**, **Extended Data Fig. 6e**, **Supplementary Video 8**). In accordance, immunostaining against PDZD8, a mitochondria-ER tethering protein^42^, revealed a higher fluorescent signal at constriction sites of centrally dividing mitochondria (**Extended Data Fig. 6f, g**). This conditional involvement of ER contacts in midzone fissions lends insight to previous reports that most, but not all fissions engage ER contacts (60-90%)^11,43^ or include mtDNA replication (77%)^31^. Similarly, we found that actin consistently polymerized on constriction sites of midzone, but not peripheral divisions (**Fig. 5c, d**). SiRNA-based knock-down of Inf2, a formin protein accelerating actin polymerization at the ER^38^, led to decreased rates of midzone fissions, while having no effect on peripheral fission rates (**Fig. 5e**, **Extended Data Fig. 7a**).

**Fig. 5.**
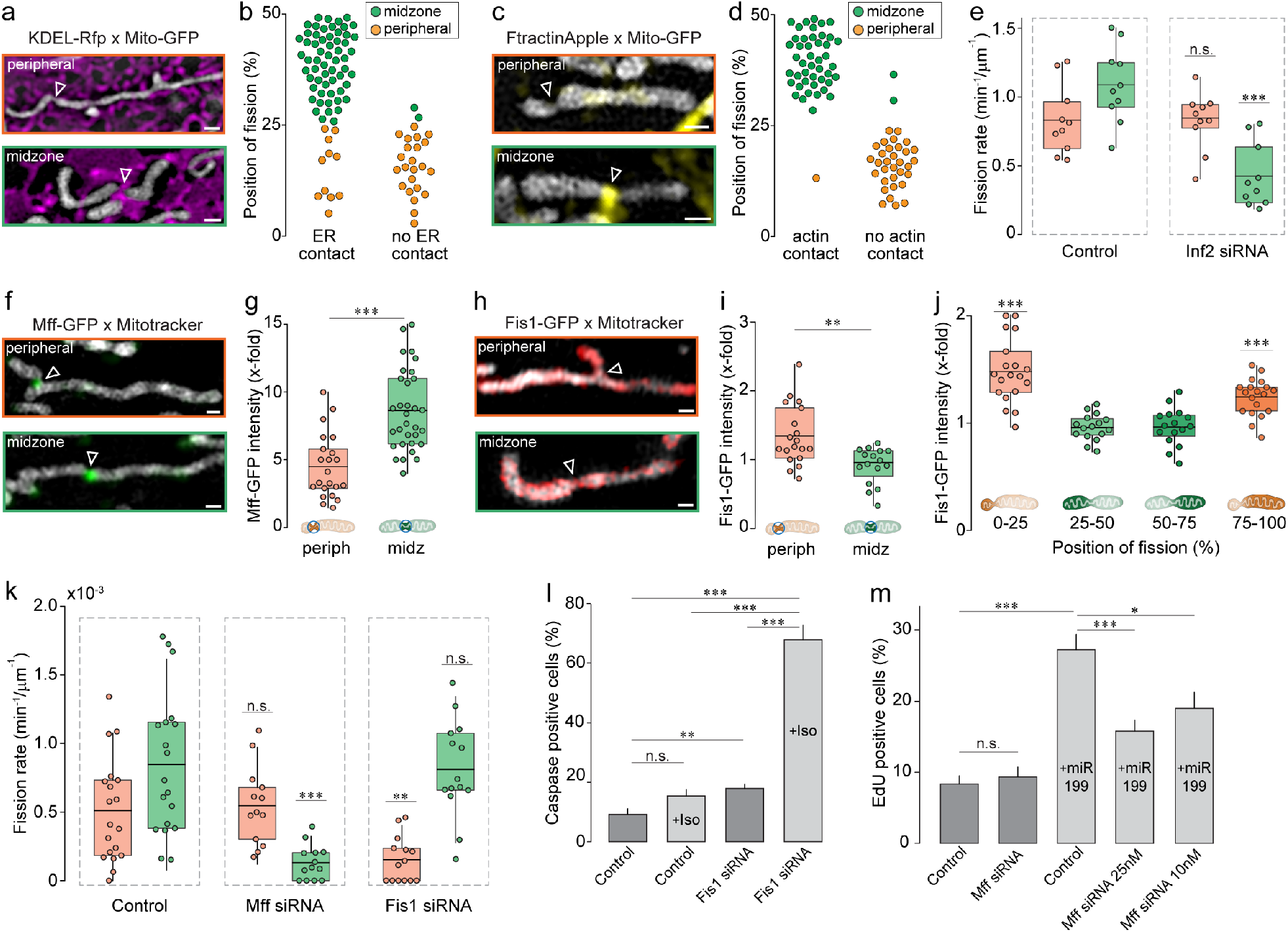
Peripheral and midzone fissions involve different molecular machineries. **a**, Two-color SIM images of mitochondria (mito-GFP, grey) and ER (KDEL-RFP, magenta) before peripheral (top) and midzone (bottom) fissions. **b**, Relative position of fission in dividing mitochondria that show a contact (left) or no contact (right) with the ER prior to fission (orange dots: peripheral, green dots: midzone; n= 86 fissions) in Cos-7 cells. **c**, SIM images of Cos-7 mitochondria (mito-GFP, grey) and actin (Ftractin-Apple, yellow) before peripheral (top) and midzone (bottom) fissions. **d**, Relative position of fission in dividing mitochondria that show a contact (left) or no contact (right) with actin prior to fission (orange dots: peripheral, green dots: midzone; n= 80 fissions). **e**, Rate of peripheral and midzone fissions in control cells versus cells treated with Inf2 siRNA (n≥ 10 FOV for each group). **f**, SIM images of U2OS cells endogenously expressing Mff-GFP and labelled with Mitotracker red displaying a peripheral (top) and midzone (bottom) fission. **g**, Normalized Mff-GFP intensity at the fission site of peripheral (left) and midzone (right) divisions (orange dots: peripheral, green dots: midzone; n= 54 fissions). Box plots display the 25/75 quartile and average (black line) with whiskers extending to 1.5 IQR. **h**, SIM images of U2OS cells endogenously expressing Fis1-GFP and labelled with Mitotracker red displaying a peripheral (top) and midzone (bottom) fission. **i**, Normalized Fis1-GFP intensity at the fission site of peripheral (left) and midzone (right) divisions (orange dots: peripheral, green dots: midzone; n= 54 fissions). **j**, Normalized Fis1-GFP intensity at the entire mitochondrial surface of the smaller and larger daughter mitochondrion of peripheral and midzone divisions measured before fission (n= 35 fissions). **k**, Quantification of the peripheral (orange) and midzone (green) fission rates in control Cos-7 cells and cells treated with Mff-siRNA or Fis1-siRNA (n≥ 13 cells for each group). **l**, Percentage of apoptotic (Caspase 3/7 positive) cardiomyocytes in control and Fis1 depleted cells after stimulated with Isoproterenol or without (n> 218 cells for each group). **m**, Percentage of proliferating (EdU positive) cardiomyocytes in control and Mff depleted cells after treatment with miR-199 and untreated (n≥ 1540 cells for each group). Statistical significance calculated by two-tailed t-test for normally distributed populations and Mann Whitney U test for non-normally distributed populations; n.s. P>0.05, *P<0.05, **P<0.01, ***P<0.001. Scale bars are 0.5 μm. Constriction sites are indicated by white arrowheads.

Finally, different adaptors are known to recruit Drp1 to outer mitochondrial membranes, but whether their roles are redundant or distinct remains unclear. To investigate their roles in different types of fission, we generated U2OS cell lines expressing endogenously tagged Mff-GFP or Fis1-GFP by Crispr/Cas9 technology. Dual-color live-cell SIM revealed strong differences in the distribution of Mff and Fis1. Mff forms bright foci at the constriction site of midzone fission, and to a lesser degree also on peripheral fissions (**Fig. 5f, g**, **Extended Data Fig. 6h, i**). In contrast, Fis1 decorates the outer mitochondrial membrane more evenly without accumulating at constriction sites. However, we observed a significant enrichment of Fis1-GFP on peripherally dividing mitochondria, compared to non-dividing and centrally dividing mitochondria (**Fig. 5h-j**), consistent with Mff and Fis1 antibody staining in fixed Cos-7 cells (**Extended Data Fig. 6h-k**). We did not find differences in accumulation of Mid49/51, which also act as Drp1 adaptors^44^ (**Extended Data Fig. 6l**). To further explore the distinct roles of Mff and Fis1, we quantified the rates of peripheral and midzone fissions in siRNA-treated cells (**Fig. 5k**, **Extended Data Fig. 7b**). Knockdown of Mff decreased midzone fission rates, while peripheral fission rates remained unchanged compared to untreated control samples. Conversely, knockdown of Fis1 reduced peripheral fission rates without affecting midzone fissions.

To examine the implications of adaptor depletions, we studied their impact on cardiomyocyte response to perturbations. We induced contraction for 48 h in cardiomyocytes and subsequently quantified apoptosis in control and Fis1-depleted cells (**Fig. 5l**). Indeed, 68% of stimulated cardiomyocytes lacking Fis1 underwent apoptosis (in contrast to 9-18% in control groups), underlining the importance of peripheral fission for cell survival under stress conditions. Finally, we knocked down Mff in cardiomyocytes that were stimulated to proliferate. While the control group showed increased cell proliferation upon miR-199 treatment, proliferation rates were significantly reduced in cardiomyocytes depleted of Mff in a dose-dependent manner (**Fig. 5m**). This further highlights the necessity of Mff-mediated midzone fissions for mitochondrial biogenesis during cell proliferation.

## Discussion

Previous studies reveal that fission is important for both mitochondrial proliferation and degradation^3^. We discovered organelle-level regulation that reconciles this paradox, with the positioning of the fission site as a key morphological determinant of the decision to proliferate or degrade. The formation of small daughter mitochondria to sequester damaged components has an advantage: the mitochondrion becomes small enough to be engulfed by autophagosomes, allowing the mass of decomposed mitochondrial material to be minimized. Our fine spatiotemporal analyses of mitochondrial bioenergetic readouts revealed that the majority of peripheral divisions are preceded by decreased membrane potential and proton motive force (matrix pH) as well as elevated ROS and Ca^2+^ levels, well before the molecular constriction of mitochondrial membranes by Drp1. While it is remarkable that a gradient forms within one mitochondrion, biological systems of all sizes frequently use gradients for monitoring their size or orientation^45^, and even individual cristae form independent units that can have different membrane potentials^46^. Thus, we speculate that peripheral fissions could use this gradient as a positioning cue, in the absence of an ER contact to define the location of the division site. This would explain how cells may regulate peripheral and midzone mitochondrial division independently, namely by an intrinsic regulation depending on mitochondrial physiological state. Independent regulation is further implied by our observation that Drp1 exploits different outer membrane adapters to perform fission: Mff recruits Drp1 to midzone fissions while Fis1 recruits Drp1 to peripheral fissions. Such a paradigm can account for previously reported variability and seeming redundancy in the machinery at division sites^47^. Our observation that Mff accumulates at peripheral fission sites, although to a lesser degree, may be driven by curvature, since we also see it enriched at the highly curved mitochondrial poles. This is consistent with reports of Mff accumulation at constricted mitochondrial regions independent of Drp1 presence^48^. We also observe lysosomes recruited to peripheral fission sites, the mechanism may be through Fis1 which is reported to recruit the lysosome-mitochondria tethering molecule TBC1D15 to the outer mitochondrial membrane^33^.

The existence of two mechanistically and functionally distinct fission types has implications in the context of pathology or aging. An excessive fission rate has been observed as a hallmark of diverse diseases^49^, and its pharmacological inhibition has been proposed as potential therapy^50–52^. Other therapeutic approaches have aimed to stimulate mitochondrial biogenesis^53,54^. However, approaches that inhibit all mitochondrial fissions might produce undesirable side effects. Our models suggest that if under pathological conditions only one type of fission is dysregulated, treatment with global inhibitors may further disrupt cell homeostasis. Therefore, our findings point towards more rational and specific therapeutic targets.

## METHODS

No statistical methods were used to predetermine sample size. For studies involving multiple experimental conditions, studies were performed on cells originated from the same cell line batch and randomly assigned to experimental conditions.

### Plasmids and reagents

Mito-GFP (Cox-8 presequence) was a gift from Hari Shroff (NIH, Bethesda), mCherry-Drp1 and BFP-KDEL were gifts from Gia Voeltz (Addgene, plasmid #49152 and #49150)^1^, SypHer mt was a gift from Nicolas Demaurex (Addgene plasmid #48251)^2^, pLPCX mito Grx1-roGFP2 was a gift from Tobias Dick (Addgene plasmid #64977)^3^, pCMV Cepia3mt was a gift from Masamitsu Iino (Addgene plasmid #58219)^4^, Lamp1-GFP was a gift from Ron Vale (Addgene plasmid #16290)^5^, YFP-Parkin was a gift from Richard Youle (Addgene plasmid #23955)^6^ and EGFP-LC3 was a gift from Karla Kirkegaard (Addgene plasmid #11546)^7^, CMV-mito-R-GECO1 was a gift from Robert Campbell (Addgene plasmid #46021^8^. FASTKD2-eGFP and mito-Twinkle-eGFP were gifts from Jean-Claude Martinou. Ftractin-Apple was a gift from Henry N. Higgs (Dartmouth College, Hanover), Mito-tagRFP was amplified from Mito-GFP.

The stable FASTKD2-eGFP cell line was recently described^9^. In brief, co-transfection of HEK293T cells with pWPT_FASTKD2-eGFP and packaging plasmids pMD2.G and psPAX2 was achieved using calcium phosphate precipitation. Medium containing virus was collected 48 hours after transfection and filtered using membranes with a pore size of 0.45 μm. The viral supernatant and polybrene were added to 70% confluent recipient cells. Culture medium was then replaced 24 hours after infection. FACS sorting was performed to select for cells expressing GFP.

Oligonucleotides for siRNA were made by Microsynth to knock down Mff (sense strand 5’-CGC UGA CCU GGA ACA AGG A-dTdT-3′), Fis1 (sense strand 5′-CGA GCU GGU GUC UGU GGA G- dTdT-3) and INF2 (sense strand 5’-GCA GUA CCG CUU CAG CAU UGU CAT T-3’ and 5’-GGA UCA ACC UGG AGA UCA UCC GCT T-3’). siRNA transfection was performed using Lipofectamine RNAi Max (Invitrogen) and cells were imaged 72 h after transfection.

The following reagents were also used: Mitotracker Green ((ThermoFisher M7514), Mitotracker Red FM ((ThermoFisher M22425), Tetramethylrhodamine ethyl ester perchlorate (Sigma, 87917), MitoSOX™ Red Mitochondrial Superoxide Indicator (Thermo Fisher M36008), Mitoquinol 98% (Adipogen SA CAY-89950-10), Quant-iT PicoGreen (Life Technologies P7581), CellEvent Caspase-3/7 (Life Technologies C10723), Fis1 polyclonal antibody (Proteintech 109561-AP), Mff (C2orf33) polyclonal antibody (Life Technologies PA567357), PDZD8 polyclonal antibody (Life Technologies PA553368), Mid49 (SMCR7) polyclonal antibody (Life Technologies, PA559950), TOM20 mouse monoclonal antibody (Santa Cruz sc-17764) and Alexa fluorophore-conjugated secondary antibodies from Life Technologies.

### Generation of knock-in U2OS cells

Crispr/Cas9-mediated knock-in of EGFP into the MFF locus directly upstream of the start codon was performed using a plasmid encoding EGFP 2 flanked by 729 bp of upstream and 723 bp of downstream homologous DNA sequence cloned into pEX-A258 to create plasmid pTW343 (pEX-A258- eGFP_hMFF_Template KI). Two pairs of sgDNA targeting MFF (sgDNA1: forward 5’- aaacAGTGATGTGTCACTGCTTGTC-3’ and reverse 5’ CACCGACAAGCAGTGACACATCACT-3’, sgDNA2: forward 5’- CACCGCATTTAAATACAGTAAATAC and reverse 5’- aaacGTATTTACTGTATTTAAATGC-3’) were cloned into pSpCas9n(BB)-2A-Puro (PX462) V2.0 (a gift from Feng Zhang (Addgene plasmid # 62987)^10^.

Crispr/Cas9-mediated knockin of EGFP into the FIS1 locus directly upstream of the start codon was performed using a plasmid encoding EGFP 2 flanked by 721 bp of upstream and 624 bp of downstream homologous DNA sequence cloned into pEX-A258 to create plasmid pTW343 (pEX-A258- eGFP_hFIS1_Template KI). Two pairs of sgDNA targeting FIS1 (sgDNA1: forward 5’- CACCGCTGAACGAGCTGGTGTCTG-3’ and reverse 5’ aaacCAGACACCAGCTCGTTCAGC-3’, sgDNA2: forward 5’- CACCGCTCGTTCAGCACGGCCTCCA and reverse 5’- aaacTGGAGGCCGTGCTGAACGAGC-3’) were cloned into pSpCas9n(BB)-2A-Puro (PX462) V2.0 (a gift from Feng Zhang (Addgene plasmid # 62987). To knock in eGFP into either the MFF or FIS1 loci, U2OS cells were co-transfected with pSpCas9n(BB)-2A-Puro (PX462) V2.0 constructs carrying the appropriate sgDNA sequences and linearized Template plasmids (pTW343 for hMFF and pTW344 for hFIS1) using Lipofectamine 2000. 24 hours after transfection, GFP-positive cells were individually isolated by fluorescence-activated cell sorting. Clones were expanded and were validated by PCR genotyping of genomic DNA and fluorescent imaging.

### Cell culture and transfection

Cos-7 cells were grown in Dulbecco’s modified Eagle medium (DMEM) supplemented with 10% fetal bovine serum and maintained in culture for a maximum of 20 passages. 10-24 h prior to transfection, cells were plated on 25 mm glass cover slips (Menzel, #1.5) at 1 × 10^5^ cells ml^−1^. Plasmid transfections were performed with 1.5 μl Lipofectamine 2000 (Invitrogen) per 100 μL Opti-MEM media (Invitrogen). The following amounts of DNA were transfected per ml: 150 ng Mito-GFP, 250 ng Mito-tagRFP, 100 ng Drp1-mCherry, 300 ng Tom20-RFP, 400 ng Mito-SypHer, 400 ng Grx1-roGFP2, 150 ng Mito-R-Geco, 400 ng Cepia3-mt, 350 ng KDEL-RFP, 250 ng, Lamp1-GFP, 350 ng, EGFP-LC3, 400 ng YFP-Parkin and 200 ng MitoTwinkle-GFP. Cells were imaged the next day.

### Primary mouse cardiomyocyte culture

Cardiac myocytes were prepared from ventricles of P1 neonatal mice using Pierce Cardiomyocyte Isolation Kit (Life Technologies 88281) following manufacturer’s instructions. After digestion, the adherent non-myocyte cells were removed by pre-plating in 10cm tissue culture plates for 45 minutes. The non-adherent cardiomyocytes were seeded in complete Cardiomyocyte plating medium supplemented with 10% fetal calf serum and antibiotics at a density of 4-5 × 105 cells/well on Poly-L-Lysine (Sigma P-7890) and Gelatin (Sigma G9891)-coated coverslips in 6-well plates. To stimulate contraction, 24h post-plating the cardiomyocytes were fed with fresh medium supplemented with 2% FCS and with 10-5 M Isoproterenol (Sigma I-2760). To induce proliferation, cardiomyocytes were transfected with 50 nM miR-199 mimic (Pharmacon, miRIDIAN Mimic Has-miR-199a-3p; C300535-05) using Lipofectamine RNAi-Max following manufacturer’s instructions (Life Technologies, 13778). Cardiomyocytes were observed 48 post-treatment or transfection.

### Western blotting

For immunoblots 48-72 hours after siRNA transfection, Cos-7 cells were lysed in RIPA buffer (Sigma) supplemented with fresh proteases inhibitors (Sigma Aldrich 11836170001) on ice for 30 minutes. A centrifugation at 16,000 g for 10 min at 4°C was performed to remove the insoluble material. Protein concentrations were determined using a Pierce BCA protein assay kit (Life Technologies, 23227) and equal amounts of protein were analyzed by self-casted 7.5 or 15% SDS-PAGE (30-50 μg of proteins per lane). For immunoblotting, proteins were transferred to nitrocellulose membranes (BioRad) electrophoretically and incubated with the specified primary antibodies (see above), diluted in 5% non-fat dry milk in Tris buffered saline with Tween 20 (TBST). The blots were further incubated with anti-rabbit or anti-mouse HRP conjugated secondary antibodies (GE Healthcare) and visualized using ECL (GE Healthcare). Where required, images of Western blotting were treated for contrast enhancement and densitometric analyses were performed using ImageJ.

### Primary antibodies used for Western blot

The following primary antibodies were used for Western blots: anti-Fis1 (LuBioScience GmbH, 10956-1-AP, diluted 1:2000), anti-Mff (Life Technologies, PA5-52765, diluted 1:500-1:1000), anti-Inf2 (Sigma-Aldrich HPA000724, diluted 1:2000), anti-alpha-Tubulin (Santa Cruz sc-5286, diluted 1:2000).

### Live-cell treatments

*TMRE:* Cells were incubated with 500 nM TMRE for 10 minutes followed by rinsing in PBS.

*Mitotracker:* Cells were incubated in 500 nM Mitotracker for 5 minutes followed by rinsing in PBS.

*PicoGreen:* to image mtDNA, cells were stained with PicoGreen diluted 1:500 for 20 min.

*MitoSox:* Cells were incubated with 5uM MitoSox for 2-4 hours prior to imaging.

*MitoQ:* Cells were incubated with 500nM MitoQ for 30-45 minutes prior and during imaging following the manufacturer’s instructions and a fresh aliquots were made every day.

### Bromouridine tagging of RNA

Cos-7 cells were incubated with 5 mM 5-bromouridine (BrU) in complete culture medium for 1 h before fixation, as described previously {Jourdain et al 2013}. BrU was stored at −20 °C, and heated and vortexed before use. Samples were immunolabeled with anti-bromodeoxyuridine (BrdU) (Roche 11170376001; 1:250 to 1:500 dilution) to visualize BrU signal.

### SIM imaging

Single and dual-color SIM imaging was performed on an 3D NSIM Nikon inverted fluorescence microscope (Eclipse Ti; Nikon) equipped with an electron charge coupled device camera (iXon3 897; Andor Technologies). The microscope was equipped with a 100x 1.49 NA oil immersion objective (CFI Apochromat TIRF 100XC Oil; Nikon). Live-cell imaging was performed at 37°C using a 488 and 561 nm laser. Acquisition setting were adapted to yield the best image quality with minimal photobleaching (laser power 0.5-15%, Exposure time 30-100 ms). Images were captured using NIS elements software (Nikon) at temporal resolution of 1 s for single-color and 6-8 s for dual-color imaging. Imaging was performed at 37°C in pre-warmed Leibovitz medium. Each sample was imaged for a maximum of 90 minutes.

### Instant SIM (iSIM)

Single and dual color instant SIM imaging was performed on a custom-built microscope setup as previously described {York et al 2013; Curd et al 2015}. Fluorescence was collected with a 1.49 NA oil immersion objective (APONXOTIRF; Olympus), with 488 nm and 561 nm excitation lasers and an sCMOS camera (PrimeBSI, 01-PRIME-BSI-R-M-16-C, Photometric). Images were captured at 0.5-5 s temporal resolution for both channels. All imaging was performed at 37°C in Leibovitz media. Raw iSIM images were subsequently deconvolved using the Lucy-Richardson deconvolution algorithm^11^ provided by Hari Shroff implemented in MATLAB, and were run for 40 iterations.

### Confocal imaging

Ratio-metric imaging of Grx1-rGFP was performed on an inverted microscope (DMI 6000; Leica) equipped with hybrid photon counting detectors (HyD; Leica). The sample was excited sequentially frame by frame with 408 nm and 488 nm with the detection set to 500-535 nm. Fluorescence was collected through a 63x 1.40 NA oil immersion objective (HC PL APO 63x/1.40 Oil CS2; Leica). Images were captured using the LAS X software (Leica). All imaging was performed at 37°C in pre-warmed Leibovitz medium for maximum 90 min per sample.

For confocal microscopy of fixed BrU samples the imaging was performed using a Leica TCS SP8 inverted microscope equipped with 405-, 488-, 552- and 638-nm lasers and a Plan-Apochromat oil objective (×63, NA 1.4). The Lightning mode (Leica) to generate deconvolved images. Microscope acquisitions were controlled by LAS X (v. 3.5.2) software from Leica.

CLEM and Caspase 3/7 samples were imaged on a Zeiss LSM 700 inverted confocal microscope with equipped with a Plan-Apochromat oil objective (×63, NA 1.40) and 488-nm and 555-nm solid-state lasers and three photomultipliers. Acquisitions were controlled by the Zeiss Zen (v. 6.0.0) software.

### Immunofluorescence

Cells were seeded on glass cover slips and grown to a confluence of ~80%. Fixation of cultured cells was performed in cold 4% paraformaldehyde (PFA) in phosphate-buffer saline (PBS) for 20 min, then cells were washed 3x in PBS. Subsequently cells were incubated with 0.3% Triton X-100 and 1% pre-immune goat serum for 30 min. The same buffer was used to incubate cells with the specified primary antibody (see above) over night at 4°C. After washing in PBS, cells were incubated with the appropriate secondary antibody for 1h and rinsed in PBS before imaging.

### Immunohistochemistry on cardiomyocytes

CMs (P1) were transfected with 25nM siFis1or siMff for 6 hours and then transfected with 50 nM of Mir-199 mimic for 48h. EdU was added in fresh medium after 56 hours and kept for the last 18 hours. Cells were then fixed for 10 minutes in in 4% paraformaldehyde in PBS and permeabilized with 0.3% Triton X100 in PBS. After blocking (PBS containing 0.001% Triton X100, 1% BSA and 1% FCS), cells were incubated overnight at 4 °C with anti-TroponinI (1:500). The day after, cells were washed 3 times and incubated 1 h at RT in the dark with the secondary-conjugated antibody diluted 1:500 (488 goat anti-rabbit A11008 Life Technology). EdU has been labelled and detected using a Click-iT EdU Alexa Fluor 594 Imaging Kit (Invitrogen C10339). Nuclei were stained with DAPI (Invitrogen).

### Correlated confocal and TEM

Cells were seeded on gridded coverslips (MatTek, P35-1.5-14-CGRD-D) and grown to 50-60% confluence. Cells were fixed at room temperature for 1 h in freshly prepared fixative (2% PFA, 1% glutaraldehyde in PBS 0.1M, pH 7.4), followed by 10x washing in PBS. Samples were imaged by confocal microscopy on the same day and z-stacks were acquired of whole cells, the pinhole was closed to 0.5 AU and pixel size reduced to 50-100 nm in xy and 100-150 nm in z. Samples were stored overnight, in PBS at 4°C. They were then stained with osmium and potassium ferrocyanide, followed by osmium alone, each with cacodylate buffer. They were finally stained with 1% uranyl acetate, then washed in water, dehydrated through 15 increasing concentrations of alcohol, and infiltrated with Epon resin. This was polymerized over night at 65°C. Serial, ultra-thin serial sections were then cut of the cell of interest, and the sections collected on single slot copper grids with a formvar support membrane. Images were recorded in a transmission electron microscope operating at 80kV (FEI Company, Tecnai Spirit).

### Image analysis

All image analysis was performed with the open-source ImageJ/Fiji^12,13^ (including Weka Segmentation, EMBL bleach correction plugins). Mitochondrial fissions were defined as events, where a single mitochondrion divided into two independently moving daughter mitochondria in live-cells and in fixed cells when a (Drp1-positive) constrictions site showed a diameter of <180 nm, measured via FWHM across the constriction. For representation purpose, a 1 pixel Gaussian Filter was used and some videos where bleach corrected using bleach correction.

#### Relative position of constriction site

The positioning of constriction site was measured manually by drawing a line along the length axis of the mitochondrion in the frame before fission. For branched mitochondria (~13 % of mitochondrial population), the length of individual branches were summed.

#### Relative fluorescent intensity

Intensity measurements of biosensors were analyzed by measuring mean fluorescence intensity on each side of the constriction and in single daughter mitochondria after fission (ROI defined by using Otsu thresholding) on SIM images and subtracting the cytosolic background. For normalization, the mean fluorescence intensity was measured in three non-dividing mitochondria within the same field of view at the same time-point. For fixed samples (anti-PDZD8, anti-Mff, anit-Fis1, anti-Mid49), the mean intensity was measured in a 500 nm circle placed at the constriction site and normalized over the fluorescent signal along the non-constricted part of the mitochondrion. For measuring Drp1, Lamp1-GFP, Mff-GFP and Fis1-GFP intensities, a 500 nm circle was placed at the constriction site and normalized over the intensity of at a 500 nm circle placed on a non-constricted part of the same mitochondrion.

#### Fission rate

To measure the fission rate, the total mitochondrial volume was calculated using trainable Weka segmentation (Fiji plugin) followed by binarization of the image. The total mitochondrial length was calculated using the total volume in the binarized image divided by the mean mitochondrial diameter. Fission rates were indicated as number of fissions occurring per μm length of mitochondria, per minute.

#### ER or actin contacts

A contact between mitochondria and ER or actin was measured by placing a line along the length axis of the mitochondrion crossing the constriction site and measuring the intensity profile for both channels. A contact site was defined, if the two signals cross at the constriction site (mitochondrial signal drops and ER signal increases at least 2x over background) for at least three consecutive frames before fission.

#### Lysosome-mitochondria contact

A contact between mitochondria and lysosome was categorized as a close proximity (<500 nm) between lysosomes and the mitochondrial constriction site for at least three consecutive frames before fission.

#### BrU quantification

Individual cells were selected manually, using Fiji’s rectangular selection tool. Three consecutive slices to focus at the bottom of the cell were then chosen upon inspection & foci were detected automatically in both BrU, and FASTKD2 channels using a fixed threshold. Foci that occurred in both channels were then counted as relevant mitochondrial RNA-transciption granules, whereas nuclear transcription for instance, was excluded from the analysis. Multiple thresholds were tried, where 150 a.u. provided the most sensible results in control cells and are represented in Extended Data Fig. 3g, though all other data are provided in the source-data files, and followed the same trend. The same pipeline was applied to both Control and UV-treated samples. The relevant Fiji-macro scripts & metadata allow for 100% traceability of the results, and will be available at github.com/TimoHenry or upon reasonable request.

### Statistics

Sample sizes were determined on the basis of prior experience in previous experiments. All statistics were performed using OriginPro software. First, datasets were tested for normal distribution using D’Agostino-Pearson normality test (significance value of 0.05). If a dataset failed this test, a non-parametric test was chosen to compare significance of means between groups (Mann-Whitney test for two samples and Kruskal-Wallis test for more than two samples). For normally distributed datasets, a t-test was chosen to compare two samples and ANOVA was used for more than two samples. Critical comparative datasets (Fis1 and Mff datasets, Drp1 intensity analysis, ER analysis) were partially re-analyzed or analyzed with an automated analysis pipeline to exclude observer bias. p values < 0.05 were considered to be significant and indicated by “*”; p values < 0.01 were indicated by “**” and <0.001 by “***.”

### Data and materials availability

All imaging as well as numerical data relevant to this study are or will be publicly available on the online repository Zenodo (doi 10.5281/zenodo.3550643) or upon reasonable request.

## ACKNOWLEDGMENTS

We thank Cécilia Cottiny and Mariona Colomer for experimental and technical assistance, Mary-Claude Croisier and Graham Knott from the BioEM (EPFL) for carrying out the electron microscopy and Thierry Laroche and the BIOP (EPFL) for imaging support. We thank Thomas Misgeld, Jean-Claude Martinou and Pavan Ramdya for feedback on the manuscript. Research in S.M.’s lab is supported by the National Centre of Competence in Research Chemical Biology, the European Research Council (CoG 819823, Piko) and the Swiss National Science Foundation (182429). T.K. received funding from European Molecular Biology Organization (ALTF-739-2016) and the Munich Cluster for Systems Neurology. This work is supported by grants from the Swiss National Science Foundation to T.P. (no CRSII5_173738 and no 31003A_182322).

## AUTHOR CONTRIBUTIONS

T.K. and S.M. conceived the project and designed experiments. T.K. performed imaging experiments and analysis. T.R. performed the Mito-Twinkle, FASTKD2, BrU and MitoSox imaging and contributed to the analysis. T.K. and J.W. performed the Caspase, Drp1 and LC3 imaging. D.M. developed and adjusted the iSIM setup. T.K., T.R. and D.M. performed the TEM experiments. S.Z. performed the western blots. F.R., M.N. and T.P. designed and performed culturing of mouse cardiomyocytes and the proliferation assay. T.W., S.Z. and H.P.L. designed and cloned the Crispr/Cas9 transgenic lines. T.K. and S.M. designed figures and wrote the manuscript, with input from all authors.

## SUPPLEMENTARY INFORMATION

Supplementary information is available (Supplementary figures 1-7, Supplementary videos 1-8).

