## Supplementary Information for "Distinct molecular signatures of fission predict mitochondrial degradation or proliferation"

Laboratory of Experimental Biophysics
École polytechnique fédérale de Lausanne
Rt de la Sorge 15
CH- 1025 Lausanne

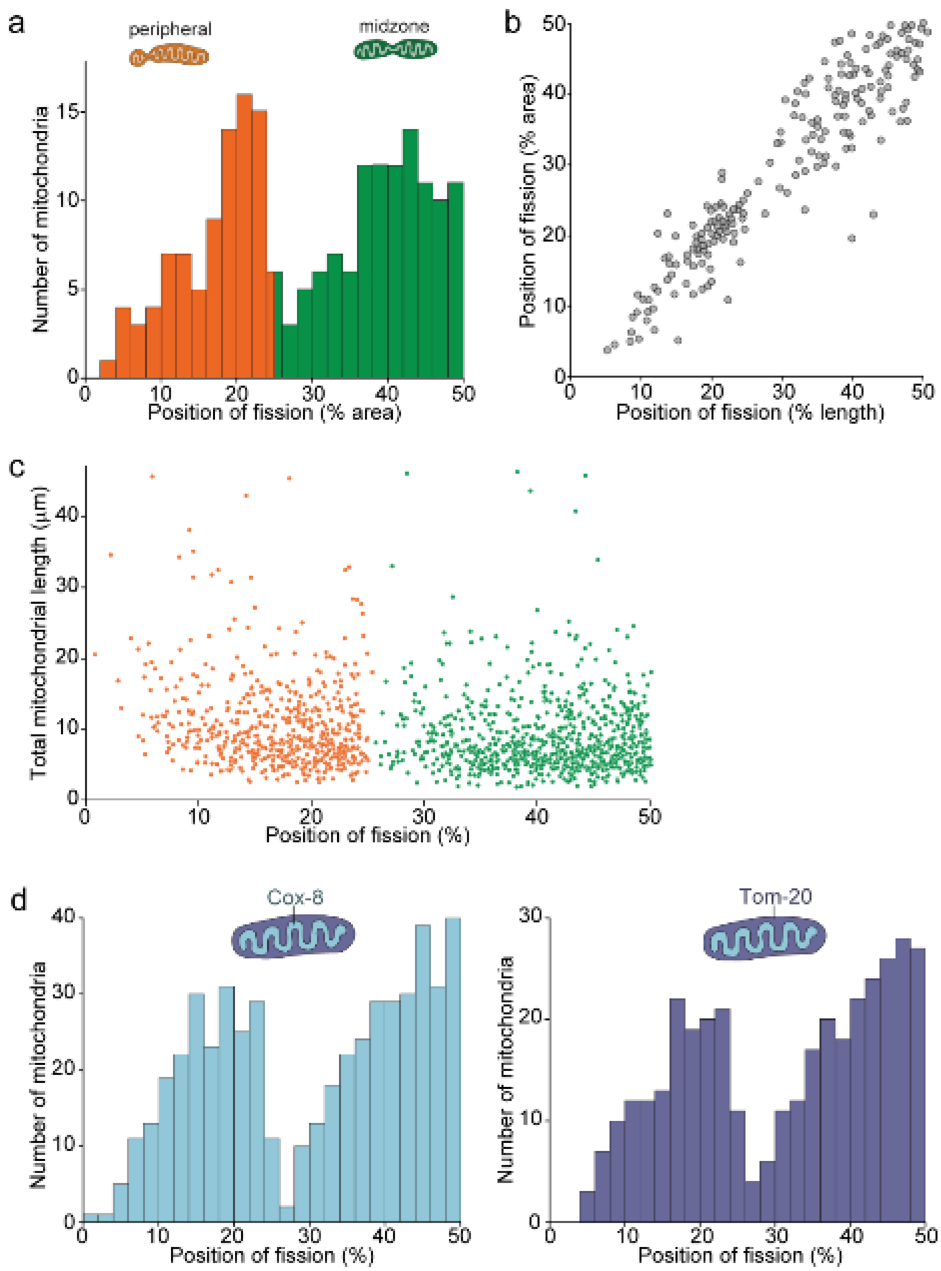


**Extended Data Fig. 1 | Distribution of mitochondrial fission sites. a**, Histogram of the relative position of constriction/fission measured by mitochondrial volume (n=201 fissions). The two peaks are colored in orange (0-25 bin; 'peripheral position') and green (25-50 bin; 'midzone position'). **b**, Relative position of mitochondrial fission measured by length versus measured by area (n=201 fissions) **c**, Scatter plot of the total length of dividing mitochondria versus the relative position of the fission site along the length axis with peripheral fissions (0-25% bin) colored in orange and midzone fissions (25-50% bin) in green (n=1393 fissions). **d**, Histogram of relative position of fission in datasets acquired with a mitochondrial inner membrane marker (left, Cox-8 targeting domain; n=510 fissions) and a mitochondrial outer membrane marker (right, TOM20; n=368 fissions).

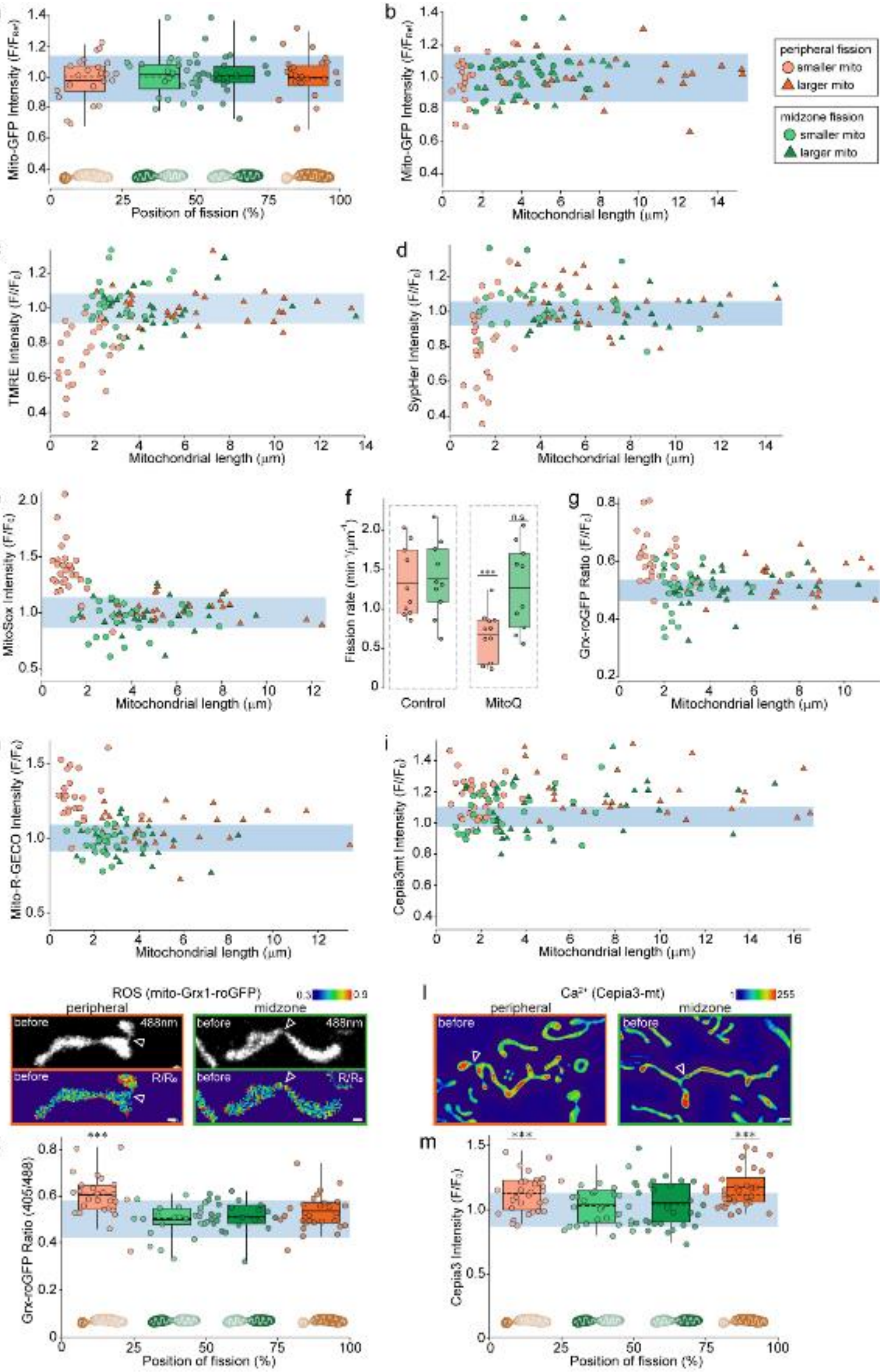
a
Mito-GFP Intensity (F/F_ref)
Position of fission (%)
b
Mitochondrial length (μm)
peripheral fission
smaller mito
larger mito
midzone fission
smaller mito
larger mito
c
TMRE Intensity (F/F₀)
d
SypHer Intensity (F/F₀)
e
MitoSox Intensity (F/F₀)
f
Fission rate (min⁻¹/μm⁻¹)
***
n.s.
Control
MitoQ
g
Grx-roGFP Ratio (F/F₀)
h
Mito-R-GECO Intensity (F/F₀)
i
Cepia3mt Intensity (F/F₀)
j
ROS (mito-Grx1-roGFP)
peripheral
midzone
before
488nm
R/R₀
l
Ca²⁺ (Cepia3-mt)
k
Grx-roGFP Ratio (405/488)
m
Cepia3 Intensity (F/F₀)

**Extended Data Fig. 2 | Physiological changes preceeding fission is independent of volume and absolute length.** **a**, Normalized mito-GFP intensity depending on the relative position of fission measured in mitochondria immediately before fission. Circles indicate individual measurements and average values of binned groups (0-25%, 25-50%, 50-75% and 75-100%;) are represented as box plots (box= 25-75 percentile; bars= min/max values; dotted line=median; solid line= average; n≥ 23 fissions for each group). Light blue area shows average mito-GFP intensity in non-dividing mitochondria (±SD). **b**, Dependence on the length of the daughter mitochondria of normalized Mito-GFP intensity immediately before peripheral (orange) or midzone (green) fissions (n≥ 23 for each group). Light blue area shows the average mito-GFP intensity in non-dividing mitochondria (±SD) **c**, Dependence on the length of the daughter mitochondria of normalized TMRE intensity immediately before peripheral (orange) or midzone (green) fissions (n≥ 28 for each group). Light blue area shows the average TMRE intensity in non-dividing mitochondria (±SD). (**d**) Dependence on the length of the daughter mitochondria of normalized mito-SypHer intensity before fission (n≥ 25 for each group) represented as in **b**. **e**, Dependence of the length of the daughter mitochondria on normalized MitoSox intensity before fission (n≥ 24 for each group), represented as in **b**. (**f**) Rates of peripheral and midzone fissions in control Cos-7 cells versus cells treated with 500nM of the ROS scavenger MitoQ (n= 10 FOVs for each group). (**g**) Dependence on the length of the daughter mitochondria of ratio-metric intensity of Grx-roGFP immediately before fission (n≥ 25 for each group) represented as in **b**. **h**, Dependence on the length of the daughter mitochondria of normalized mito-R-Geco intensity before fission (n≥ 31 for each group) represented as in **b**. (**i**) Dependence on the length of the daughter mitochondria of normalized Cepia3-mt intensity before fission (n≥ 31 for each group) represented as in **b.** **j**, Examples of Grx1-roGFP transfected mitochondria dividing in the periphery (left) and midzone (right) taken from ratiometric measurements. **k**, Ratio-metric intensity of Grx-roGFP depending on the position of fission represented as in **b.** Light blue area shows the average Grx-roGFP ratio in non-dividing mitochondria (±SD; n≥ 25 fissions for each group. **l**, Images of Cepia3-mt transfected mitochondria immediately before peripheral (left) and midzone (right) fission. **m**, Normalized Cepia3-mt intensity depending on the position of fission, represented as in **b**. Light blue area shows the average Cepia3-mt intensity in non-dividing mitochondria (±SD; n≥ 31 fissions for each group). Statistical significance calculated by two-tailed t-test for normally distributed populations and Mann Whitney U test for non-normally distributed populations; ns > 0.05, ***P<0.001. Scale bars are 0.5 µm. Fission sites are indicated by arrowheads.

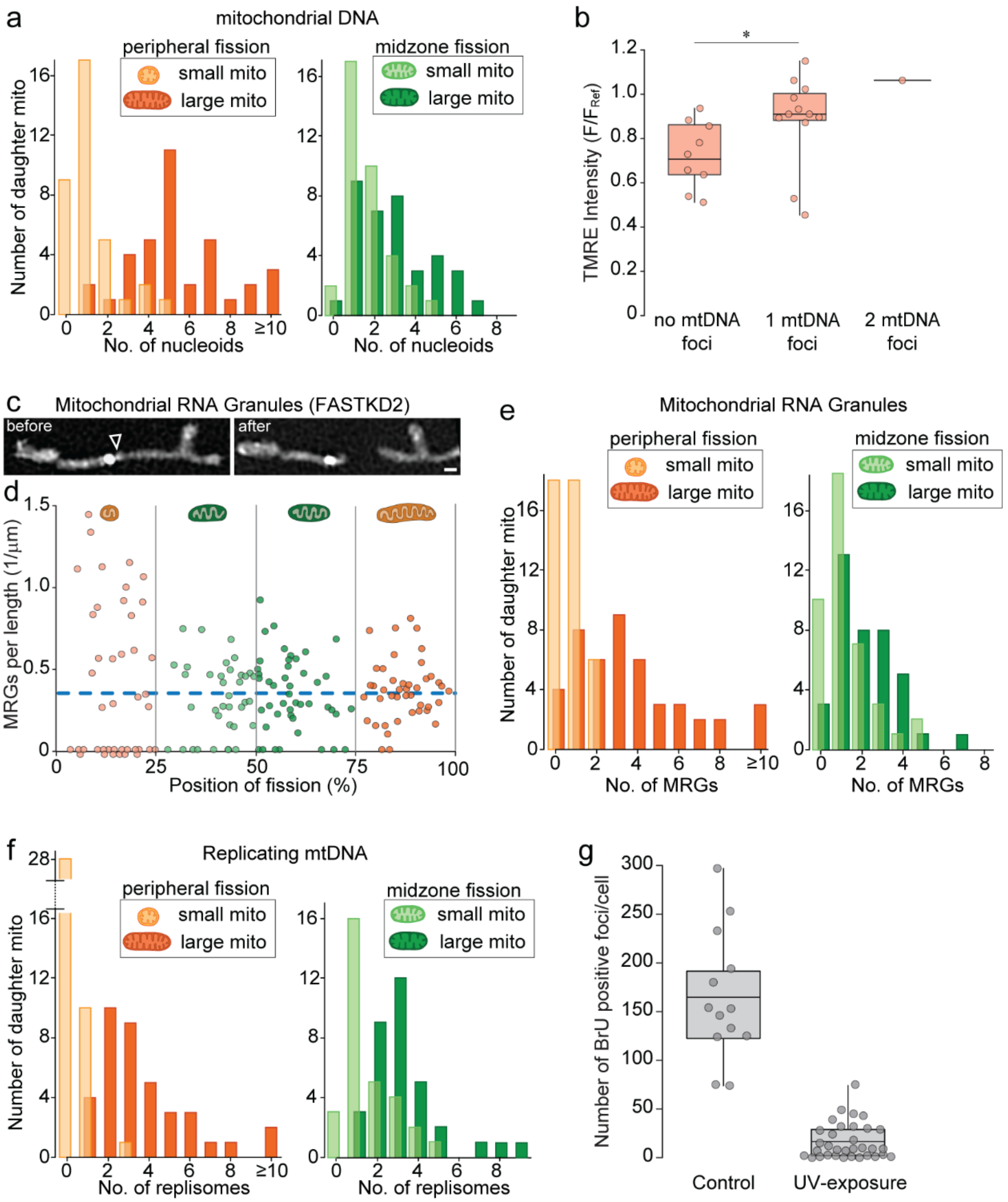


**Extended Data Fig. 3 | Redistribution of mitochondrial DNA and RNA granules.** **a**, Distribution of of PicoGreen foci in the small and large daughter mitochondrion derived from peripheral and midzone fission (n≥ 35 fissions for each group). **b**, Normalized TMRE intensity in the small daughter mitochondria from peripheral fissions that contain 0, 1 or 2 nucleoids (n= 21 fissions; box= 25-75 percentile; bars= min/max values; solid line= average). **c**, SIM images of mitochondrial RNA granules (MRGs, FASTKD2) before and after fission. **d**, Number of MRGs per μm length as a function of fission position. Circles indicate the individual measurements color coded for binned groups (0-25%, 25-50%, 50-75% and 75-100%; n≥ 35 for each group). Blue line shows average MRG per length in non-dividing mitochondria (n=41) **e**, Distribution of the number of MRGs (FASTKD2; n≥ 41 fissions for each group) and **f**, replicating nucleoids (mito-Twinkle; n≥ 34 fissions for each group) in smaller (light) and larger (dark) daughter mitochondria from peripheral (left, orange) and midzone (right, green). **g**, Number of BrU positive foci per cell in control Cos-7 cells and cells exposed to UV light for 3 min prior to measurement (n≥ 39 cells for each group). Statistical significance calculated by two-tailed t-test for normally distributed populations and Mann Whitney U test for non-normally distributed populations; *P< 0.05, ***P<0.001. Scale bar is 0.5 μm.

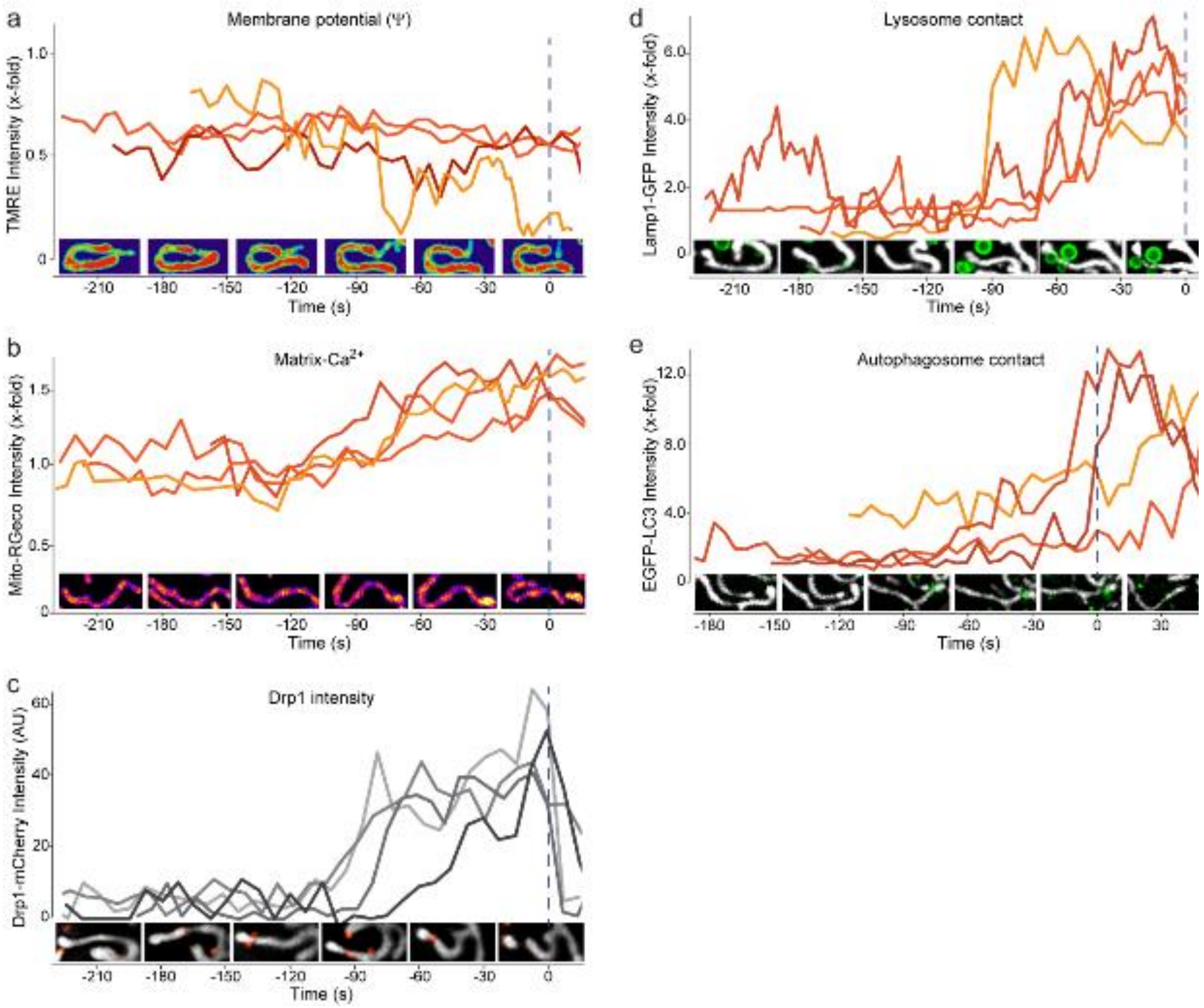


**Extended Data Fig. 4 | Time-course of physiological changes and recruitment of fission regulators.** Time course of fluorescent signals in four examples of Cos-7 mitochondria displaying normalized **a**, TMRE intensity, **b**, mito-R-Geco intensity and **c**, Drp1 intensity with corresponding SIM images in the mitochondrial compartment giving rise to the smaller daughter mitochondria before a peripheral division. **d**, Time course of lysosome co-localization and **e**, autophagosome co-localization at constriction sites for peripheral fissions, by measuring Lamp1-

mEGFP and EGFP-LC3B intensity respectively. For EGFP-LC3B measurements, cells were pre-treated with 10 μM CCCP to increase LC3 signals. Blue dotted line (t= 0s) marks the timepoint of fission.

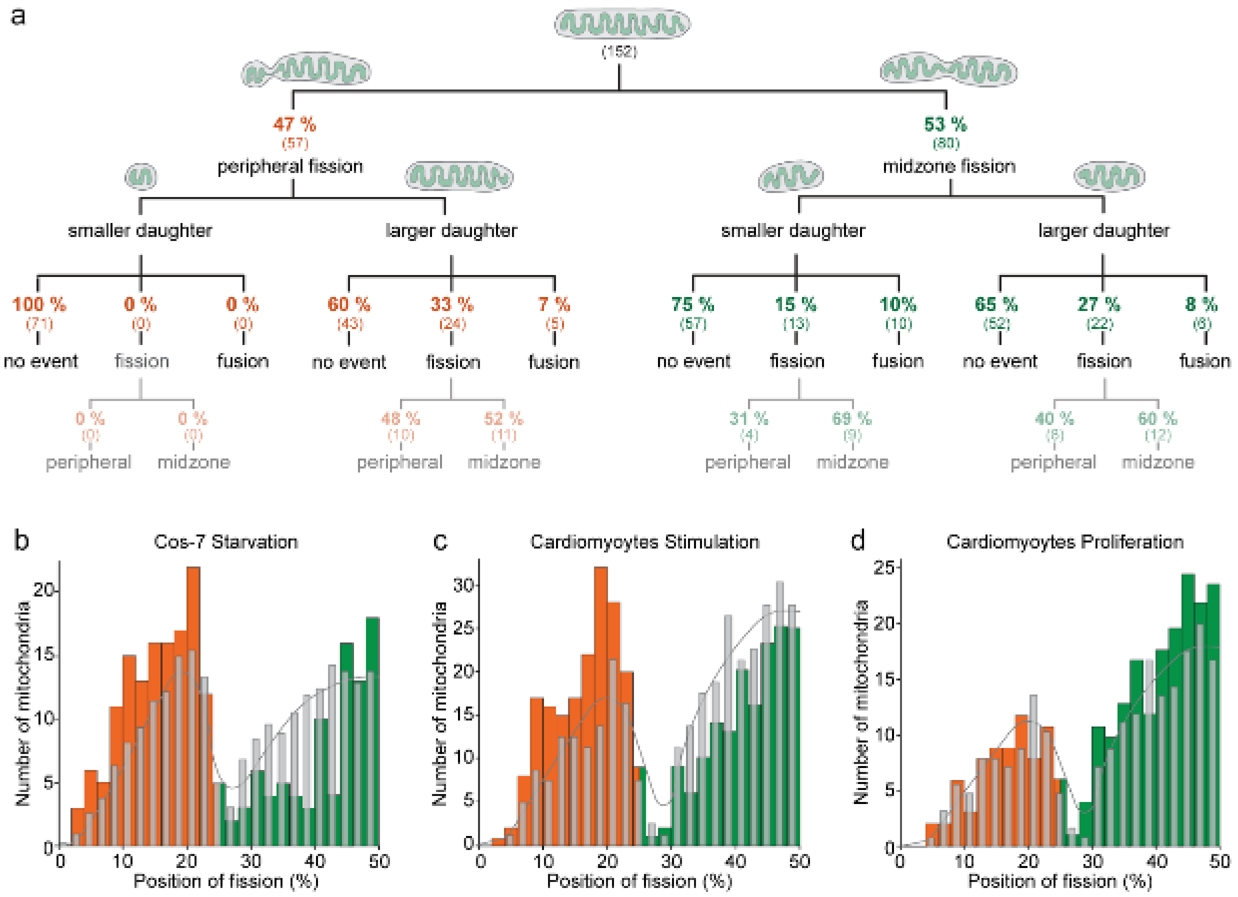


**Extended Data Fig. 5 | Peripheral and midzone fissions interact differently with the mitochondrial network and the distribution of the fission positions is regulated independently. a**, Schematic diagram depicting the fate (”no event”, another “fission” or “fusion”) of each daughter mitochondrion from peripheral and midzone fissions after the initial division in postnatal cardiomyocytes. Only mitochondria, that could be traced for more than 100 seconds post-fission were included in the analysis. **b**, Distribution of the relative position of in starved Cos-7 cells, with peripheral (1-25%) fission labeled in orange and midzone fissions (25-50%) in green (n=212 fissions). The frequency distribution of Cos-7 control samples is superimposed in grey (replotted from **Fig. 1c**). **c**, Distribution of the relative position of constriction/fission along the length axis of isoproterenol treated mouse cardiomyocyte mitochondria (n=356 fissions) and **d**, miR-199 treated cardiomyocytes (n=223 fissions) respectively. The frequency distribution of untreated mouse cardiomyocytes samples is superimposed in grey (replotted from **Fig. 1g**).

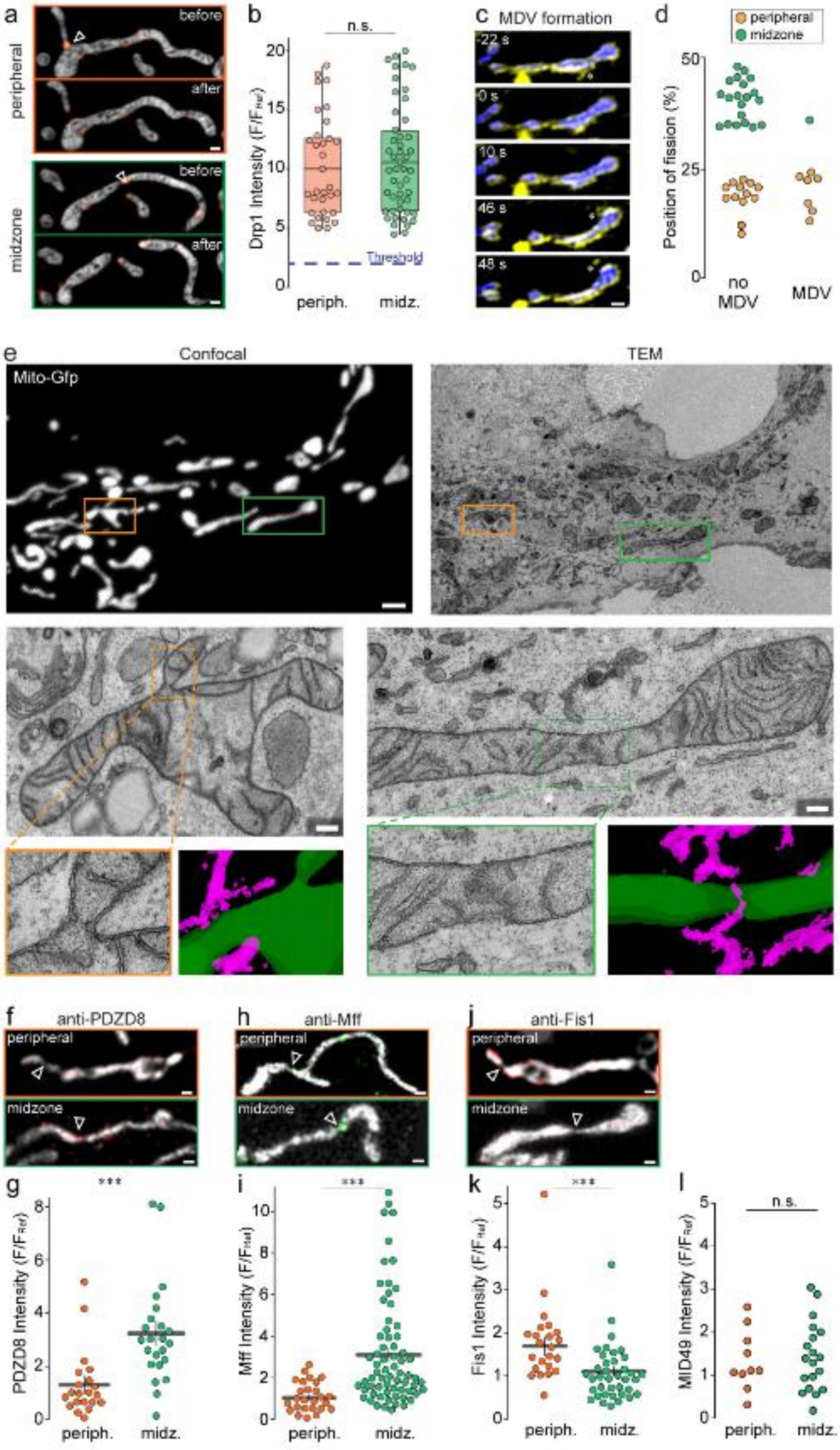

a
peripheral
before
after
midzone
before
after
b
n.s.
Drp1 Intensity (F/F_Ref)
Threshold
periph.
midz.
c
MDV formation
-22 s
0 s
10 s
46 s
48 s
d
peripheral
midzone
Position of fission (%)
no MDV
MDV
e
Confocal
TEM
Mito-Gfp
f
anti-PDZD8
peripheral
midzone
h
anti-Mff
peripheral
midzone
j
anti-Fis1
peripheral
midzone
g
***
PDZD8 Intensity (F/F_Ref)
periph.
midz.
i
***
Mff Intensity (F/F_Ref)
periph.
midz.
k
***
Fis1 Intensity (F/F_Ref)
periph.
midz.
l
n.s.
MID49 Intensity (F/F_Ref)
periph.
midz.

**Extended Data Fig. 6 | Peripheral and midzone fissions are both Drp1 mediated but involve distinct upstream mechanisms.** **a**, Two-color SIM images of mitochondria (mito-GFP, greyscale) and Drp1 (Drp1-mCherry, red) undergoing peripheral or midzone fission. **b**, Normalized Drp1 intensity on the constriction sites of peripheral and midzone divisions. The threshold for a Drp1 accumulation (blue dotted line) was set at a signal > 3x over background (n≥ 48 for each group). **c**, Time-lapse sequence of a SIM movie, where both mitochondrial outer membrane (TOM20-GFP) and inner membrane (mito-RFP) were labeled to detect MDV formation (asterisks). **d**, Quantification of the fission positions for mitochondria undergoing MDV formation or not before or after division (n= 41 fissions). **e**, Correlated confocal and transmission electron microscopy (CLEM) of mitochondria in Cos-7 cells labeled with Mito-GFP, fixed 24 h after expression. Zoom in of two individual mitochondria with a peripheral (orange frame) and a midzone (green frame) constriction. A pseudo-coloring of three consecutive TEM z-sections recombined into a single rendering shows mitochondria (green) and ER (magenta). Scale bar represents 2 μm in confocal and 200 nm in TEM images. **f**, SIM images of asymmetric and symmetric constrictions in fixed Cos7 cells labeled with anti-Tom20 (grey) and anti-PDZD8 (red), **h**, anti-Mff (green) and **j**, anti-Fis1 (red). (**g**) Distribution of normalized fluorescent intensities of anti-PDZD8 (n= 48 fissions), **i**, anti-Mff (n= 91fissions) and **k**, anti-Fis1 (n= 59 fissions) staining in fixed Cos7 cells for asymmetric (orange) and symmetric (green) fissions. **l**, Distribution of normalized fluorescence intensities of anti-MID49 (n≥ 48 for each group) staining in fixed Cos7 cells for peripheral (orange) or midzone (green) fissions. Statistical significance calculated by two-tailed t-test for normally distributed populations and Mann Whitney U test for non-normally distributed populations; n.s. P>0.05, ***P<0.001. Scale bars are 0.5 μm. Fission sites are indicated by white arrowheads.

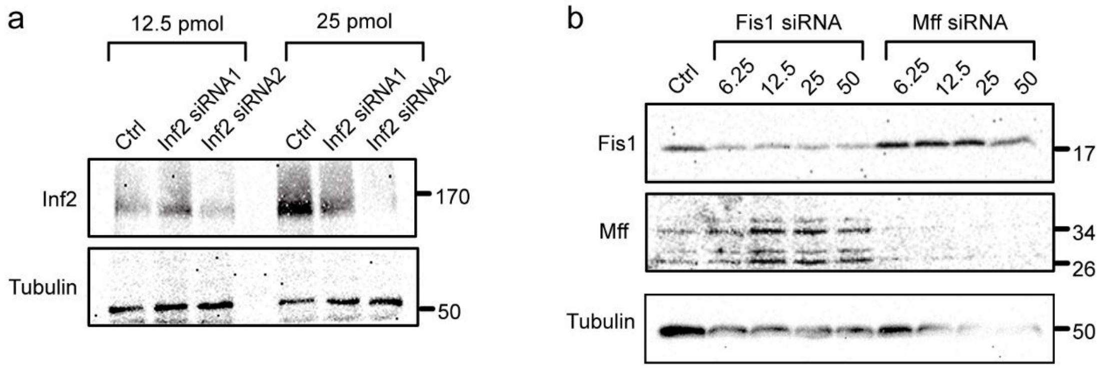


**Extended Data Fig. 7 | Silencing efficiency of Fis1, Mff and Inf2 siRNA in Cos-7 cells.** **a**, Western blot analysis of protein levels for Cos-7 cells 72 hours after transfection with two siRNAs against Inf2 at two different concentrations (pmol). Molecular mass is in kilodalton. **b**, Western blot analysis of protein levels for Cos-7 cells 72 hours after transfection with four different concentrations (pmol) of siRNA against Fis1 and Mff.

### SUPPLEMENTARY VIDEOS

**Supplementary video 1: Live imaging of mitochondrial fissions**

Live-cell SIM imaging of peripheral and midzone mitochondrial divisions in Cos-7 cells labeled with Mitotracker green. Video was acquired at 1 frame/sec for 3 minutes and corresponds to **Fig. 1 b**.

**Supplementary video 2: Live imaging of mouse cardiomyocyte**

Live-cell iSIM imaging of a post-natal mouse cardiomyocyte labeled with Mitotracker green showing peripheral (orange arrowhead) and midzone (green arrowhead) fissions. Video was acquired at 1 frame/5 sec for 6 minutes. Video corresponds to **Fig. 1 f**.

**Supplementary video 3: Live imaging of mitochondrial membrane potential (TMRE)**

Live-cell SIM imaging of TMRE stained Cos-7 cells showing a peripheral mitochondrial divisions. Video was acquired at 1 frame/3 sec. To highlight differences in fluorescent intensities, a heat-map look-up table was chosen.

**Supplementary video 4: Live imaging of mitochondrial matrix pH (Mito-SypHer)**

Live-cell SIM imaging of Cos-7 transfected with mito-SypHer showing a peripheral mitochondrial divisions. Video was acquired at 1 frame/1.5 sec. To highlight differences in fluorescent intensities, a heat-map look-up table was chosen.

**Supplementary video 5: Live imaging of mitochondrial $Ca^{2+}$ (Mito-R-Geco)**

Live-cell SIM imaging of Cos-7 transfected with mito-R-Geco showing a peripheral mitochondrial divisions. Video was acquired at 1 frame/3 sec. To highlight differences in fluorescent intensities, a heat-map look-up table was chosen.

**Supplementary video 6: Live imaging of mitochondria-lysosome contact during fission**

Live-cell iSIM imaging of lysosomes (Lamp1-mEGFP, green) and mitochondria (Mito-tagRFP, grey) in Cos-7 cells. During peripheral fission (left) but not during midzone fission (right), lysosomes contact the mitochondrial constriction site. Video was acquired at 1 frame/3 sec for 3 minutes. Video corresponds to **Fig. 3 d**.

**Supplementary video 7: Live imaging of mitochondria-autophagosome contact after fission**

Live-cell iSIM imaging of Cos-7 cells where autophagosomes (LC3-GFP, green) and mitochondria (Mito-tagRFP, grey) are labeled. After peripheral fission, the small daughter mitochondrion is subsequently engulfed by an autophagosome. Video was acquired at 1 frame/5 sec for 6.5 minutes. Video corresponds to **Fig. 3 h**.

**Supplementary video 8: Live imaging of mitochondria-ER contacts during fission**

Live-cell iSIM imaging of Cos-7 cells transfected with KDEL-RFP (ER) and mito-GFP (mitochondria) showing a midzone fission (left, green arrowhead) and a peripheral fission (right, orange arrowhead). Video was acquired at 1 frame/14 sec for 5 minutes.
